# Integrating Multiplexed Imaging and Multiscale Modeling Identifies Tumor Phenotype Transformation as a Critical Component of Therapeutic T Cell Efficacy

**DOI:** 10.1101/2023.12.06.570168

**Authors:** John W. Hickey, Eran Agmon, Nina Horowitz, Matthew Lamore, John Sunwoo, Markus Covert, Garry P. Nolan

## Abstract

Cancer progression is a complex process involving interactions that unfold across molecular, cellular, and tissue scales. These multiscale interactions have been difficult to measure and to simulate. Here we integrated CODEX multiplexed tissue imaging with multiscale modeling software, to model key action points that influence the outcome of T cell therapies with cancer. The initial phenotype of therapeutic T cells influences the ability of T cells to convert tumor cells to an inflammatory, anti-proliferative phenotype. This T cell phenotype could be preserved by structural reprogramming to facilitate continual tumor phenotype conversion and killing. One takeaway is that controlling the rate of cancer phenotype conversion is critical for control of tumor growth. The results suggest new design criteria and patient selection metrics for T cell therapies, call for a rethinking of T cell therapeutic implementation, and provide a foundation for synergistically integrating multiplexed imaging data with multiscale modeling of the cancer-immune interface.

## INTRODUCTION

Cancer is a complex system of interactions that unfold across molecular, cellular, and tissue scales (Fig. 1A). Adoptive T cell immunotherapy—in which patients are given T cells specific for cancer—causes a systems-level perturbation to cancer and has shown decisive clinical results in certain types of cancer but limited efficacy in solid tumors (Ahmed et al., 2015; Fesnak et al., 2016; Johnson and June, 2017; June et al., 2018; Neelapu et al., 2018; Restifo et al., 2012; Thistlethwaite et al., 2017; Yee, 2010, 2014). Indeed, much still needs to be learned about the manners by which infused cellular products cause effective therapeutic results. For instance, it is well understood that T cell phenotype matters, but how does cancer cell phenotype influence T cell therapy efficacy? Can infused T cells transform cancer phenotype, and is this related to T cell phenotype at the time of therapeutic delivery? Are there additional mechanisms for T cell phenotype maintenance related to their environment? How does the phenotype of the T cell product affect restructuring of the tumor tissue?

**Figure 1:**
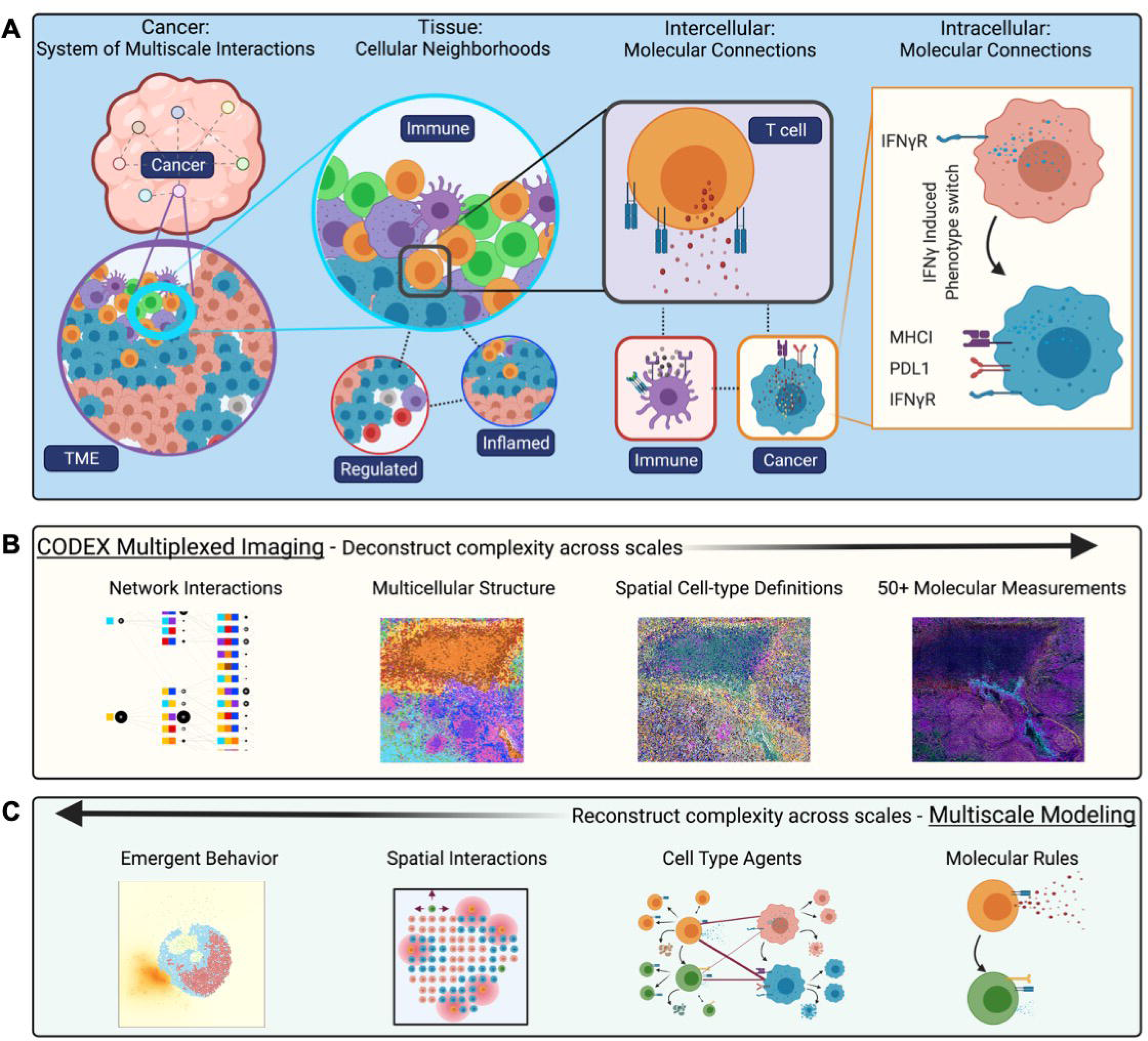
Cancer is a system of network interactions, and its analysis requires methods that can deconstruct and reconstruct the complexity at multiple scales. A) At the tissue scale, multicellular neighborhoods form to make larger tissue structures and organs. At the cellular scale, cells engage with each other through intermolecular interactions, and intracellular interactions mediate cellular function. B) CODEX imaging enables multiplexed molecular measurements of 50 or more proteins that can be quantified at a single-cell level. These molecular profiles can be used to define both cell types and states. Using the spatial features of the data, multicellular structures can be identified based on conserved composition. Finally, network interactions across these scales can be interpreted to fully <u>deconstruct</u> the complexity of a tissue. C) Multiscale modeling enables <u>reconstruction</u> of complex biology across scales. Models are defined by molecular rules for cell agents that facilitate interactions. These interactions happen within a spatial microenvironment that result in emergent biological behavior. Models can guide hypothesis generation.

Such questions remain unanswered given the difficulty of interrogating the native cancer-immune state that provides sufficiently reflective biological measurements that would allow researchers to build models to simultaneously capture the multiple scales (molecular, cellular, tissue) of cancer. Single-cell measurement technologies have allowed characterization of molecular changes in intracellular processes but do not reveal the spatial features of intercellular interactions in cancer (Galletti et al., 2020; Krishna et al., 2020). On the other hand, traditional histologic approaches can capture spatial features of cancer but is limited to only a few molecular markers at once— restricting the ability to co-define cell types or cell phenotypes *in situ* (Schietinger et al., 2013; Stack et al., 2014; Thul et al., 2017). Computational modeling has been largely restricted to a single biological scale for either description or source of input data, which limits the ability of modeling to accurately predict interactions across multiple scales. Consequently, methods are needed that can provide data that both deconstructs cancer’s interaction networks at multiple scales and that allows accurate modeling and reconstruction of such networks that allows testing predictions or hypotheses to be made.

Multiplexed imaging is a recently developed technology that enables deconstructing the complexity of tissues from the top-down with spatial features preserved (Fig. 1B) (Hickey et al., 2021a). The ability to probe more than 50 markers simultaneously using the CO-Detection by indEXing (CODEX) multiplexed imaging platform makes it possible to identify molecules, cell states, cell types, and network interactions in space (Goltsev et al., 2018; Schürch et al., 2020). The concurrent development of computational systems biology approaches has facilitated quantifying and identifying key network interactions from this data such as multicellular neighborhoods (Bhate et al., 2021).

Multiscale modeling is a complementary approach for discovering critical interactions, by reconstructing the complexity of tissues from the bottom-up with computational simulations (Fig. 1C) (An et al., 2009; Makaryan et al., 2020; Norton et al., 2019; Swat et al., 2012; Talman et al., 2019; Zhang et al., 2009). *Vivarium* is a recently introduced software tool that simplifies multiscale modeling, making it possible to connect modules of diverse mechanistic models into integrative simulations that cover multiple spatial and temporal scales (Agmon et al., 2021). This enables leveraging extensive prior knowledge about relevant biological mechanisms that were measured separately, as demonstrated previously with the construction of spatial bacteria colony models (Skalnik et al., 2021).

We leveraged both CODEX multiplexed imaging and the *Vivarium* multiscale modeling software to understand the interactions of T cell therapies with cancer at multiple scales. To date, most studies have employed either top-down (deconstructing the data through analysis) (Fraietta et al., 2018; Vodnala et al., 2019) or bottom-up (reconstructing the data with mechanistic models) (Cess and Finley, 2020; Jessica and Bagheri, 2021) approaches to the study of cancer. However, there is synergy in employing both approaches simultaneously to drive discovery of a more accurate tissue representation. As demonstrated here, multiscale modeling can be used to identify key points of the system for perturbation. Furthermore, information from multiplexed imaging feeds multiscale agent-based models by providing more accurate parameter values, initial states, and update rules.

By combining multiplexed imaging and multiscale modeling, we demonstrated that both tumor and T cell phenotype are key determinants of T cell therapeutic efficacy. T cell phenotype control has been a main focus to promote T cell longevity for killing cancer, with most approaches centering on intracellular molecular perturbation of T cells (Blank et al., 2019; Gattinoni et al., 2017). Much less attention has been given to tumor phenotype. Here we observed that the conversion of tumor phenotype was a critical determinant in the control of cancer growth. Tumor phenotype conversion was dependent on a CD8+ T cell phenotype with ability to divide rapidly (memory-like) and secrete IFNγ (effector-like), suggesting this as a design criterion/goal for T cell therapies as well as a matching patient selection metric. The results suggest that integrating modeling approach with multiplexed imaging data can provide a roadmap towards such a goal.

## RESULTS

### Changing tumor phenotype to an inflammatory state enhances T cell recognition and killing

To understand the influence of tumor cell phenotype on T cell therapeutic efficacy, we evaluated how differences in tumor phenotype influenced killing by T cells *in vitro*. We incubated B16-F10 mouse melanoma cells with IFNγ overnight to induce intracellular signaling, which is known to be critical to function and phenotype of cancer cells (Hoekstra et al., 2020; Sucker et al., 2017; Thibaut et al., 2020; Williams et al., 2020). As shown by CyTOF analysis, there was a phenotype change in about half of the tumor cells—characterized by upregulation of both anti-inflammatory (PDL1) and inflammatory (H2Db) surface markers in the group treated with IFNγ (Fig. 2A).

**Figure 2:**
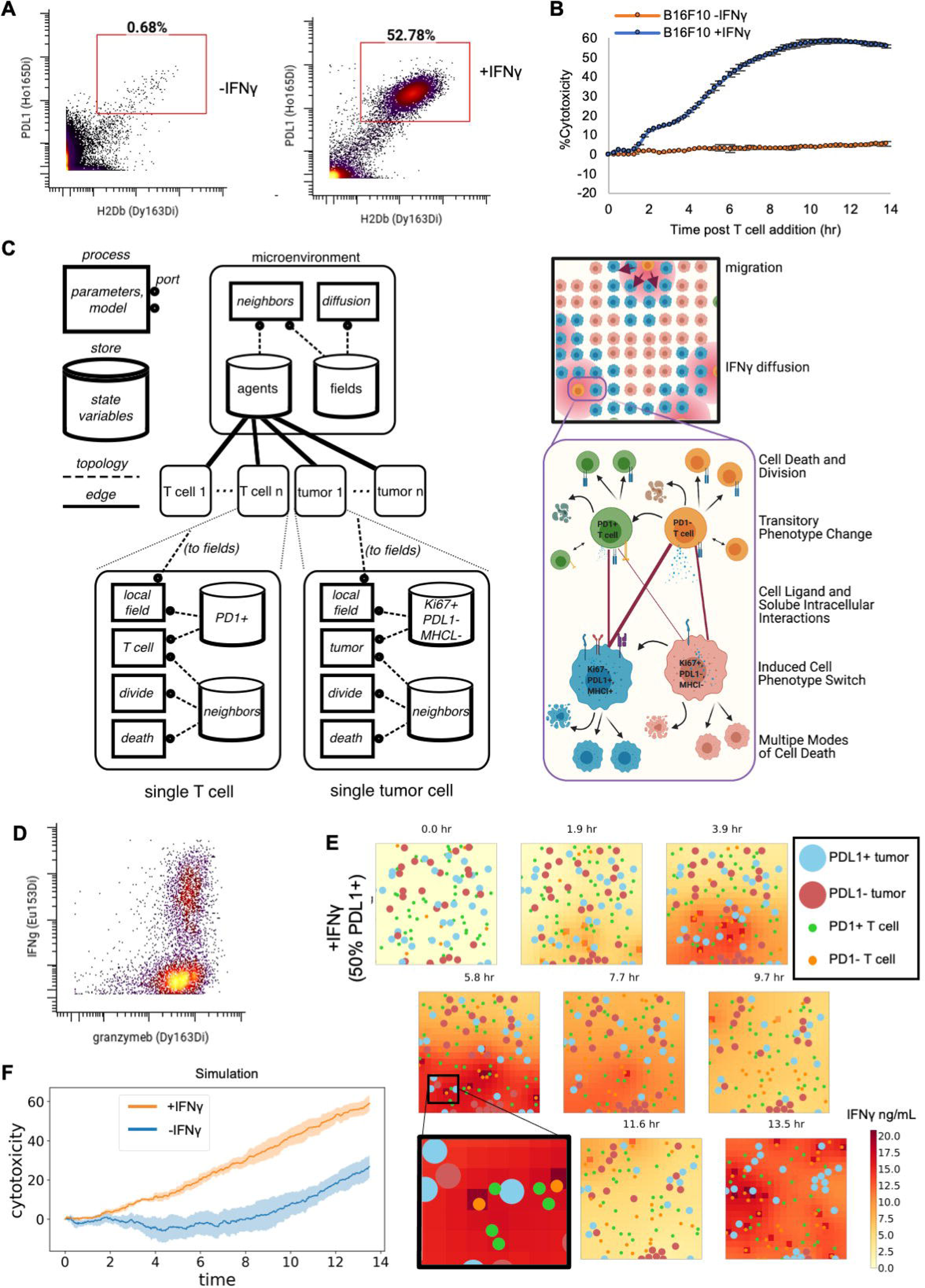
Changing tumor phenotype to an inflammatory state enhances T cell recognition and killing. **A)** PDL1 and H2Db per cell levels as measured by CyTOF of B16-F10 tumor cells after being incubated with IFNγ or no IFNγ for 18 h. **B)** Percent killing of cognate tumor cells over time by expanded therapeutic T cells pre-incubated with IFNγ or not. Tumor and T cells were incubated at a 1:1 ratio (mean of n=3 replicates with error bars showing SEM). **C)** Multiscale agent-based model of the tumor microenvironment used to understand critical components governing efficacy of adoptive T cell therapies at multiple levels of scale. **D)** Evaluation of per cell levels of effector molecules, granzymeB and IFNγ, of restimulated therapeutic PMEL CD8^+^ T cells after 10 days of activation. **E)** Snapshots of agent-based modeling results showing results from a simulation that was initialized to mirror *in vitro* killing by expanded therapeutic T cells pre-incubated with IFNγ or not. **F)** Cytotoxicity levels from multiscale agent-based modeling of initializing simulations with tumors that had similar phenotype to input tumor cells in Figure 2B indicating being treated with or without IFNγ (mean of n=5 replicates with shading showing SEM).

These tumor cells express both anti-inflammatory and inflammatory molecules, but it was unclear whether this phenotype would inhibit T cell killing or promote more efficient killing. We performed a dynamic *in vitro* killing assay with a 1:1 ratio of cognate, antigen-specific (PMEL), activated CD8^+^ T cells and B16-F10 tumor cells or IFNγ-pretreated B16-F10 tumor cells. Despite equal growth rates for the two tumor phenotypes, IFNγ-treated tumors were killed much more effectively (50% vs. 7% at 13 hours) (Fig. 2B).

To formulate a mechanistic explanation why this tumor phenotype conversion led to such enhanced killing, we created a multiscale agent-based model using *Vivarium*. The ability to create agents at multiple scales (e.g., molecular and cellular) governed by defined biological rules within a spatial environment can be used to test multiple hypotheses and detect critical inflection points in network structures and dynamics. Figure 2C illustrates how multiple biological scales of cell-state interactions is modelled with Vivarium with a simplified wiring diagram (left panel). The model was focused on interactions between two subsets of therapeutic T cells and the two subphenotypes of cancer cells. We modelled fundamental immune-tumor interactions in T cell therapy, for instance 1) PD1^+^ T cells interaction with PDL1 on the surface of PDL1^+^ MHCI^+^ tumor cells, 2) PD1^-^ T cells/PD1^+^ T cells recognition of tumor cells through interactions of the TCR with MHCI on tumor cells wherein PD1^-^ T cells can be converted to PD1^+^ T cells through repeated stimulation of their TCR and 3) CD8+ T cells release of IFNγ, which then converts tumor cells to PDL1^+^ MHCI^+^ tumor cells (Fig. 2C, right panel).

To create this model required encoding prior knowledge and lab-derived parameter values to create the rules governing individual cancer cells and T cells (see Supplemental Table 1, Fig. 1, and 2). We additionally encoded intracellular and intercellular interactions in *Vivarium* and tuned the parameters by comparing process performance with expected behavior standards*. Vivarium* also enables environmental interaction such as migration and diffusion of molecules. Jupyter notebooks explaining the development and rationale behind the model are provided on the project’s GitHub repository, where example notebooks ran different permutations of the model for testing. A documented code base also describes the rules and parameters of the model and can be found in the Methods.

We then evaluated whether our *in-silico* model would show the expected higher T cell killing efficacy of an IFNγ-induced tumor phenotype. To accurately initialize our model, we measured *in vitro* levels of PD1, effector molecules involving killing (granzyme B, perforin), and IFNγ expression from restimulated PMEL CD8^+^ T cells used for the *in vitro* killing assay (Fig. 2D, Supplemental Fig. 3). Using these values, in addition to *in vitro*-measured PDL1 and MHCI expression by tumor cells (Fig. 2A), as inputs to initialize both T cells and tumor cells in the model and ran modeling simulations that mimicked our *in vitro* killing assay setup.

To observe cellular behavior from the *T cell killing* simulation we plotted “snapshots” as output for the IFNγ-treated tumor (Fig. 2E, time of each snapshot shown above starting with the initial state a 0 h to 13.5 h). This figure thus contains rich information across spatial, time, agent, and molecular dimensions, where T cell migration, tumor proliferation, tumor phenotype change, secretion of IFNγ, and tumor killing can be observed over the course of 13.5 hours (Fig. 2E, larger circles indicate tumor cells and smaller circles indicate T cells, color represent the phenotype of a given cell type, and the red background color represents the local concentration of IFNγ). For instance, zooming in on the 5.8 h snapshot, there is a PD1+ T cell (green) interacting with a PDL1+ tumor cell (light blue) and several other tumor cells (upper left of zoomed figure) with no T cells next to them. In the next snapshot (7.7 h) this tumor cell has been killed and other tumor cells have not moved or been killed, while the T cells have all migrated. This zoomed-in area (5.8 h) also has a concentration of soluble IFNγ around 18 ng/mL with some variation in the grid squares. IFNγ has increased with the number of T cell and tumor cell interactions from a starting concentration of 0 at snapshot 0 h and decreases by the next snapshot (7.7 h) due to some T cells beginning to downregulate TCR and tumor uptake of soluble IFNγ. This led to a majority of the tumor cells changing phenotype, as can be noticed from 11.6 h to 13.5 h window (salmon color to light blue color).

We quantified the cytotoxicity in this simulation to compare with our *in vitro* data by evaluating the number of deaths and normalizing to a simulation with no T cells added. Quantification of killing by cytotoxicity indicated an important role of tumor phenotype on T cell killing even at early time points (Fig. 2F). Consequently, starting with a population of tumor cells that have a PDL1+ MHCI+ phenotype has a large impact, potentially due to increased levels of interactions between tumor cells expressing higher levels of MHCI and T cells. In summary, with identical cognate T cell inputs but different cancer cell phenotype ratio inputs, as observed *in silico* and *in vitro* data demonstrated that conversion of tumor phenotype to an inflammatory state enhances the ability of T cells to kill tumor cells, resulting in fewer total cancer cells and an inflammatory microenvironment conducive to T cell killing.

### Initial phenotype of the input therapeutic T cells drives conversion of tumor cells to an inflammatory phenotype

Controlling T cell phenotype during *ex vivo* expansion prior to therapeutic transfer is expected to be critical, especially since cells are in foreign environments for extended periods of time (Hickey et al., 2017, 2019; Kosmides et al., 2018; Oh et al., 2015). Therapeutic T cell phenotype is known to cause dramatic differences in anti-tumor efficacy, especially from the perspective of T cell persistence and effector molecule expression (Blank et al., 2019; Gattinoni et al., 2017; Henning et al., 2018; Van der Leun et al., 2020; Speiser et al., 2014, 2016). Broadly, memory T cells are expected to persist longer and give rise to more daughter cells, whereas effector cells are expected to be shorter-lived and secrete effector molecules like perforin (Farber et al., 2014; Kaech and Cui, 2012). However, because T cells are usually isolated from subjects or dissociated from cancer tissues to be measured, the manners by which their phenotype relates to tumor phenotype, at the beginning and end of therapy, remain unknown.

One approach shown to generate T cells with these distinct phenotypes, is via inhibition of acetyl-CoA production (Vodnala et al., 2019) by incubating CD8^+^ T cells with 2-hydroxycitrate (2HC) during expansion (Fig. 3A). Inhibition of acetyl-CoA formation pushes T cells toward memory stemness resulting in significantly better tumor control than conventionally activated T cells (Vodnala et al., 2019). Indeed, cells incubated *in vitro* with 2HC had lower expression of PD1^+^ than the untreated cells (25% vs. 75% PD1^+^) (Fig. 3B). Additional characterization by CyTOF showed further subphenotypes (*Hickey et al cosubmitted*), but two main categories of memory and effector T cells were broadly separated by PD1 status, and we used these as inputs to our model.

**Figure 3:**
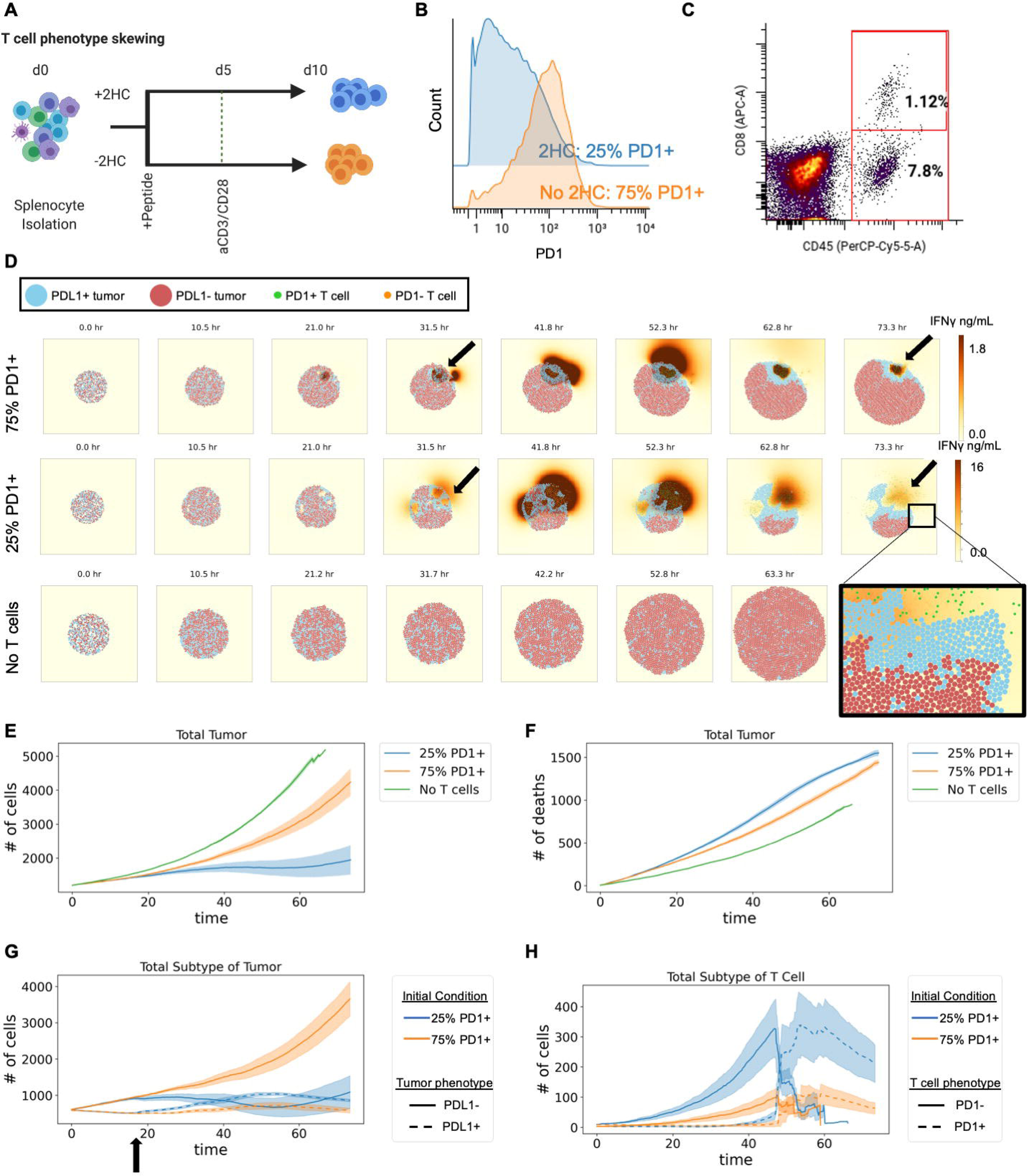
The initial phenotype of transferred therapeutic T cells influences the ability of T cells to convert tumor cells to an inflammatory phenotype. **A)** Experimental layout for controlling T cell phenotype during *ex vivo* T cell expansion. **B)** Histogram of per cell levels of PD1 expression as measured by CyTOF of T cells treated with metabolic inhibitor 2HC or without the inhibitor. **C)** Percent of CD8^+^ T cells within tumors post-treatment with therapeutically expanded T cells as measured by flow cytometry. **D)** Snapshots of simulation initialized with *in vivo*-relevant cell numbers, ratios, and T cell phenotypes for 25% and 75% PD1^+^ T cell conditions compared to a simulation condition with no T cells. **E)** Total number of tumor cells over the course of the simulation that was 3 biological days under each condition. **F)** The number of tumor cell deaths over the time course of the simulation. **G)** Number of tumor cells separated by phenotype over the course of the simulation. **H)** Number of T cells separated by phenotype over the course of the simulation. For panels E-H: mean of n=4 replicates with shading showing SEM.

We compared how these two phenotypes influence the ability of T cells to alter tumor cell phenotype under conditions found within the tumor microenvironment. For our previous *in vitro* experiments, the ratio of tumor to T cells was controlled; however, there are several barriers to entry into tumors *in vivo* (e.g., extracellular matrix, trafficking to non-tumor organ sites). Therefore, to create an accurate starting ratio of tumor to T cells in the tumor microenvironments, we transferred therapeutic T cells into mice bearing established B16-F10 tumors. This *in vivo* data showed that the CD8^+^ T cell frequency in these tumors was approximately 1% of all cells (Fig. 3C).

With biologically and therapeutically relevant initialization conditions for both phenotype and T cell ratio, we ran simulations with 1% CD8^+^ T cells (12 T cells to 1200 tumor cells) to compare our two differentially activated T cell phenotypes (with 2HC: 25% PD1^+^; without 2HC: 75% PD1^+^) as separate therapies over a period of 72 hours. Snapshots from the simulations show distinct dynamics of IFNγ secretion, CD8^+^ T cell proliferation, and tumor killing as well as different spatial phenomenon relating to tumor phenotype (Fig. 3D, Supplemental Videos 1-3). For example, tumor cells proliferated in all groups from 0 h to 31.5 h. T cells also proliferated by 31.5 h, started to kill tumor cells, and converted tumor cells to PDL1^+^ phenotype in pockets (black arrows) for both 25% and 75% PD1^+^ T cell conditions. These T cell pockets have higher local concentration of IFNγ (red/brown) than other areas of the tumor, and highest levels of IFNγ in the 25% PD1^+^ T cell condition. By 73.3 h, the T cells killed enough tumor cells to coalesce into common pockets and escape the tumor bed (magnified area, bottom right of Fig. 2D).

Quantification of the simulations showed that the starting condition of 25% PD1^+^ T cells inhibits tumor growth more effectively (∼1000 tumor cells at 60 h) than the starting condition of 75% PD1^+^ T cells (∼3000 tumor cells at 60 h) (Fig. 3E). Under conditions without T cells, the tumor cells grew exponentially (∼5000 tumor cells at 60 h). These results were expected based on previous *in vivo* experiments (Vodnala et al., 2019).

To explain the difference in tumor control *in silico*, we looked at the number of tumor cells killed in each of the conditions. Surprisingly, control of tumor growth was not explained by the number of T cell-induced killing events in the two conditions (25% PD1^+^ T cells, ∼1300 cell deaths at 60 h; 75% PD^+^ T cells, ∼1200 deaths at 60 h) (Fig. 3F, Supplemental Fig. 4A). Since tumor phenotype made a large functional difference in our *in vitro* experiments and earlier simulations, we investigated the sub-phenotypes of tumor cells in these simulations. In the 25% PD1^+^ T cell condition, there was an earlier (at 18 h indicated by black arrow) and stronger conversion of tumors from a proliferative to non-proliferating inflammatory phenotype than in the 75% PD1^+^ T cell condition (Fig. 3G). This control was linked to increased numbers of PD1-T cells in the tumor, though in both conditions T cells became PD1+ over the course of the simulation (Fig. 3H, Supplemental Fig. 4B). Thus, greater tumor control from phenotype-switched T cells was due to greater ability to inhibit tumor proliferation rather than differences in inhibition from T cell direct killing.

### T cells induce tumor cell phenotype conversion *in vivo*

Our attempts to reconstruct the complexity of the system in both *in vitro* and *in silico* models suggested the critical importance of a tumor phenotype transformation by T cells. To study this *in vivo*, we used CODEX multiplexed imaging to enable measurement of tumor phenotype changes in a spatial context in a therapeutically relevant adoptive T cell model (Fig. 4A) (Black et al., 2021; Kennedy-Darling et al., 2021; Vodnala et al., 2019). Specifically, we transferred T cells that were treated or not with 2HC into B16F10 established tumors (day 10) and harvested them 3 days after treatment (day 13) for CODEX multiplexed imaging. Imaging was performed with a 42-antibody panel designed to detect cancer phenotypic markers (e.g., PDL1, H2Kb, H2Db, Ki67), immune cell-type-defining markers (e.g., CD3, CD4, CD8, F4/80), and functional markers (e.g., PD1, CD27) (*Hickey et al. cosubmitted*).

**Figure 4:**
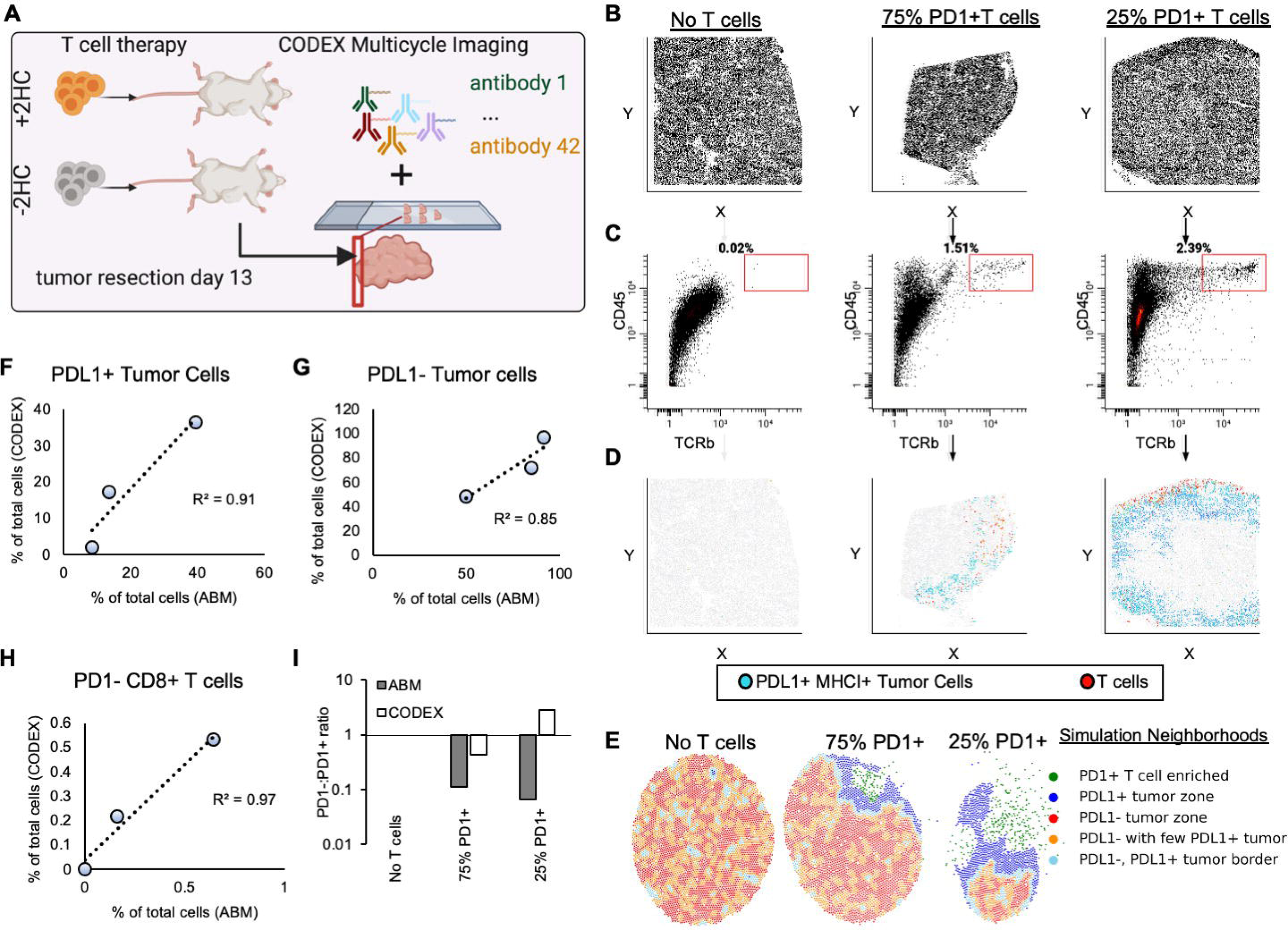
T cells induce tumor cell phenotype conversion in vivo. **A)** Experimental layout for *in vivo* adoptive T cell therapy and CODEX multiplexed imaging of tumors at day 3 post-treatment. **B-D)** CODEX multiplexed imaging results in single-cell data that is spatially resolved. Scatter plots for each treatment condition plotted for each cell in **B**) X vs. Y, **C**) CD45 vs. TCRb, and **D**) gated T cells (red) in the X vs. Y axis together with PDL1+ MHCI+ tumor cells (blue)—to see spatial distribution of T cells in the tumor samples. **E)** Multicellular neighborhood analysis for each of the simulations at the day 3 endpoint reveals differential structures created by each of the responses, where responses are characterized into 5 overall neighborhoods. **F-I)** Correlation plots between percent of cells resulting after three days of T cell therapy for both CODEX multiplexed imaging of *in vivo* experiments and *in silico* simulations for **F)** PDL1+ tumor cells, **G)** PDL1-tumor cells, and **H)** PD1-CD8+ T cells. **I)** PD1- to PD1+ T cell ratio for agent-based modeling (ABM) and *in vivo* (CODEX) experiments.

Because we observed spatial restriction of the T cells that was also associated with proximity to PDL1^+^ MHCI^+^ tumor cells within our simulations (Fig. 3D), we predicted we would see the same proximal events in our *in vivo* data. To compare results between *in vivo* and simulations we first examined the positions of T cells within tumor sections, since CODEX generates single-cell data that enables cell-type identification (Hickey et al., 2021b). Because of this we can visualize each individual cell in x and y coordinates (Fig. 4B). Each individual cell contains the quantification of each protein marker and so we used this to gate T cells and PDL1^+^ MHCI^+^ tumor cells (Fig. 4C). An analysis of the locations of T cells (TCRb^+^ CD45^+^, red) and PDL1^+^ MHCI^+^ tumor cells (blue) in each tumor section revealed spatial restriction of the T cells and co-localization with inflamed tumor cells in both T cell conditions (Fig. 4D), mirroring the Vivarium modeling prediction.

To further investigate this phenomenon, we compared the *in vivo* CODEX data and *in silico* modeling output by performing multicellular neighborhood analysis on our simulation data since both datasets preserve spatial features (Fig. 4E) (Schürch et al., 2020). In the *in silico* data, there are neighborhoods representing borders of immune attack on the tumor from both T cell-treated groups, with larger borders in tumors treated with 25% PD1^+^ T cells than 75% PD1^+^ T cells and disorganized tumor neighborhoods in tumors without T cell treatment (compare 4E middle figure to rightmost). Here, the borders were enriched in both T cells and PDL1^+^ MHCI^+^ tumor cells. These features of the *in silico* models were consistent with major structural components in the *in vivo* data (*Hickey et. al., co-submitted*), suggesting the important role of coordination of the neighborhood interactions, structure, and function. Moreover, co-localization of transformed T cells and tumor cells supports the hypothesis that T cells are responsible for tumor phenotype conversion *in vivo*.

We also compared the cell-type percentages in the CODEX data with *in silico* percentages at day 3 to understand whether relative phenotype conversion rates were similar. We found good correlations of ending percentages for PDL1^+^ tumor cells (Figure 4F, R=0.91), PDL1^-^ tumor cells (Figure 4G, R=0.85), and PD1^-^ T cells (Figure 4H, R=0.97). However, this was not the case for PD1^+^ T cells: The ratio of PD1^+^ T cells to PD1^-^ T cells was much greater in the *in silico* model than we observed in the CODEX multiplexed imaging data for the 25% PD1^+^ T cell treatment group (Figure 4I, Supplemental Fig. 5A). Because the death rate for PD1^-^ T cells was lower than for the PD1^+^ T cells in the *in silico* model, this suggests that death does not account for the ratio difference (Supplemental Fig. 5B, C). This result suggests that there is a mechanism for T cell phenotype maintenance missing from our model.

### Spatial location of T cells impacts ability to maintain phenotype

We hypothesized that discrepancies between our model and *in vivo* data provide an avenue to uncover biological mechanisms of T cell phenotype preservation by combining CODEX data with our multiscale model. Part of the reason we chose an agent-based model design is because it complements the spatial and compositional structure of CODEX multiplexed imaging. Since we quantify protein expression at the single-cell level with spatial coordinates that are linked to cell type, we can directly import our CODEX multiplexed imaging data as initial states of our multiscale simulations (Fig. 5A). This uniquely allows us to use the model to interpret complex CODEX data, extend the dynamics of static multiplexed imaging data, and establish more accurate initial conditions.

**Figure 5:**
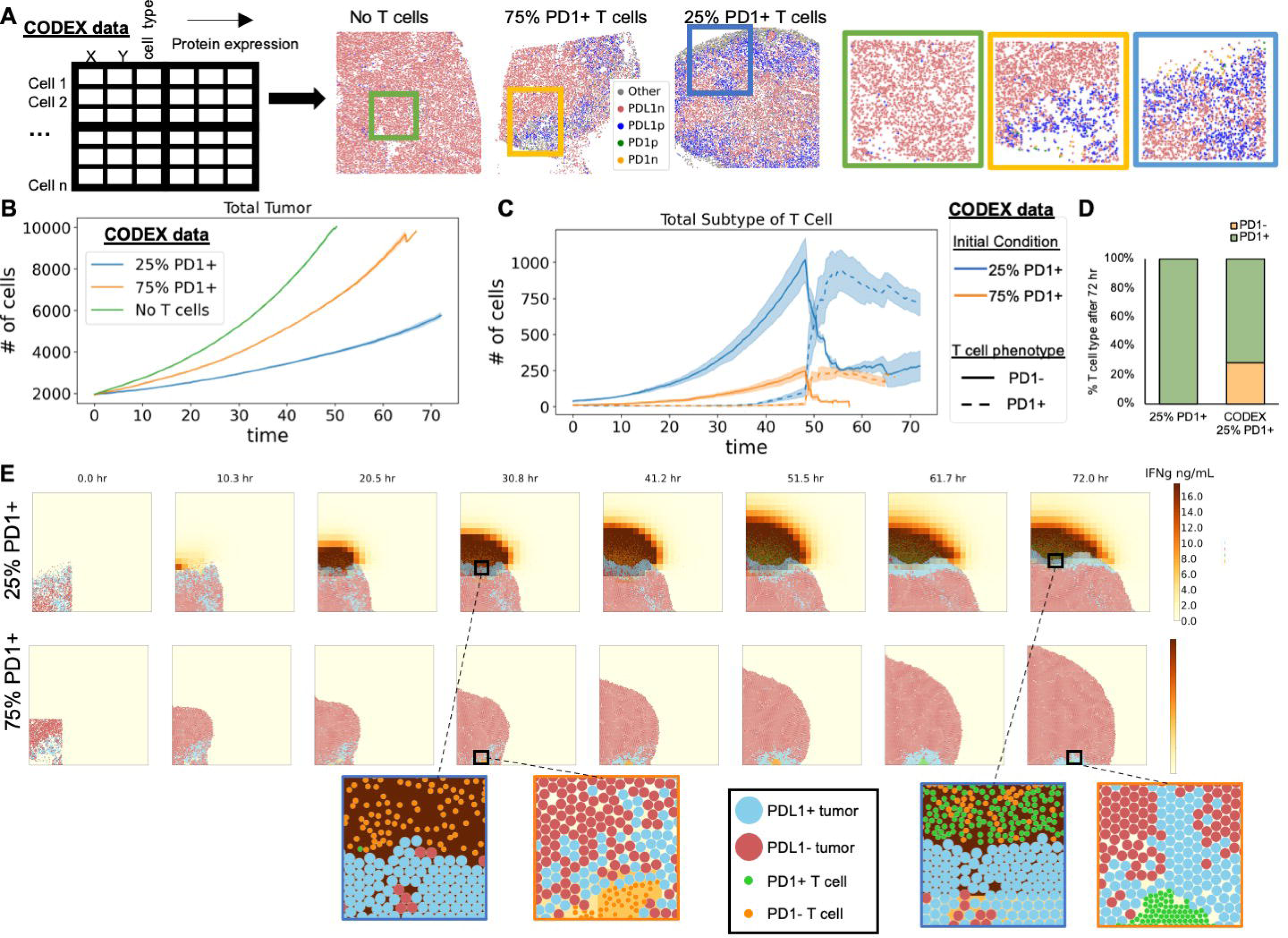
Spatial location of T cells impacts ability to maintain phenotype. **A)** Left: CODEX multiplexed data is amenable to initialize multiscale-agent based models because it has single-cell information of cell type, X & Y positions, and molecular protein expression. Middle: Cell-type maps of CODEX images of tumor sections. Rectangles indicate subsets of 2000 cells used to initialize the model. Right: High-magnification images of the areas indicated by rectangles in the middle panels. **B)** Number of tumor cells in T cell-treated and control groups as a function of simulation time (mean of n=4 replicates with shading showing SEM). **C)** Number of PD1^+^ and PD1^-^ T cells in each T cell-treated groups as a function of simulation time (mean of n=4 replicates with shading showing SEM). **D)** Percent of PD1^+^ and PD1^-^ T cells at the end of the 72-hour simulation started either with initial conditions of 25% PD1^+^ T cells (used for Fig. 3H) or the conditions based on CODEX data (used in Fig. 5C) (average of n=4 simulations for both conditions). **E)** Snapshots of the tumor from agent-based modeling condition 25% PD1^+^ T cell and 75% PD1^+^ T cells from simulations initialized with CODEX data that illustrate spatial restrictions of T cells and zoomed in regions to indicate phenotype status of T cells over time.

We were particularly interested to see if initializing our simulations with spatial information obtained from *in vivo* data would reveal the reason that the ratio of PD1^+^ T cells to PD1^-^ T cells observed in the *in-silico* model was higher than in the CODEX multiplexed imaging data (Fig. 4I). We took a region of ∼2000 cells from CODEX images to initialize our model, and then simulated the changes in the tumor microenvironment over 3 days. Tumor cell growth rates from each condition matched expectations and previous simulations (Fig. 5B, Supplemental Fig. 6A-C). Similarly, we observed an exhaustion of the T cells in both T cell treatment conditions after about 50 hours; however, only in the 75% PD1^+^ T cell-treated condition did all the T cells become exhausted, whereas in the 25% PD1^+^ T cell-treated condition a proportion of PD1^-^ T cells remained at 72 h (∼250 PD1^-^ T cells) (Fig. 5C, Supplemental Fig. 6D). This contrasts with the results of our previous simulation (Fig. 3H), where both T cell treatments led to complete phenotype conversion to PD1^+^ by the end of 3 days (Fig. 5D).

Since T cells become exhausted through chronic TCR stimulation, we hypothesized that a spatial relationship might be responsible for phenotype preservation. The snapshots from the simulations initialized with CODEX data revealed that in the *in silico* 25% PD1^+^ T cell condition there is a front of attacking T cells on the periphery of the tumor and that the tumor cells on the border with these T cells are inflamed (Fig. 5E). This can be seen by the increased IFNγ concentration at the edge of the periphery of the tumor (brown) and by zooming in on the interface of T cells and tumor cells at 30.8 h (Fig. 5E, blue square). In contrast, the T cells in the *in silico* 75% PD1^+^ T cell condition initially attacked from the periphery but were soon surrounded by proliferating cancer cells seen at 30.8 h and remained so until 72 h (Fig. 5E, orange square). This suggests that the spatial location of T cells on the periphery of tumors may be critical for T cell phenotype maintenance.

### Conversion of tumor cell phenotype is more critical for tumor control than T cell phenotype preservation

We hypothesized that since T cells were on the periphery of the tumor, they could escape chronic stimulation and thus delay exhaustion (Fig. 6A). To test this, we initialized our simulations with 25% PD1^+^ T cells conditions (as in Fig. 3), except that T cells were located outside the tumor bed rather than inside. Like our CODEX-initialized experiment (Fig. 5), we observed tumor phenotype changes on the periphery of the tumor where tumor cells contacted T cells (Fig. 6B, increase in light blue PDL1+ tumor cells). When the T cells were initialized outside the tumor, we saw a dramatic increase in total numbers of T cells compared to the numbers when T cells were initialized on the inside of the tumor (Fig. 6C). This increase resulted from a delay in T cell exhaustion (Supplemental Fig. 7A, B).

**Figure 6:**
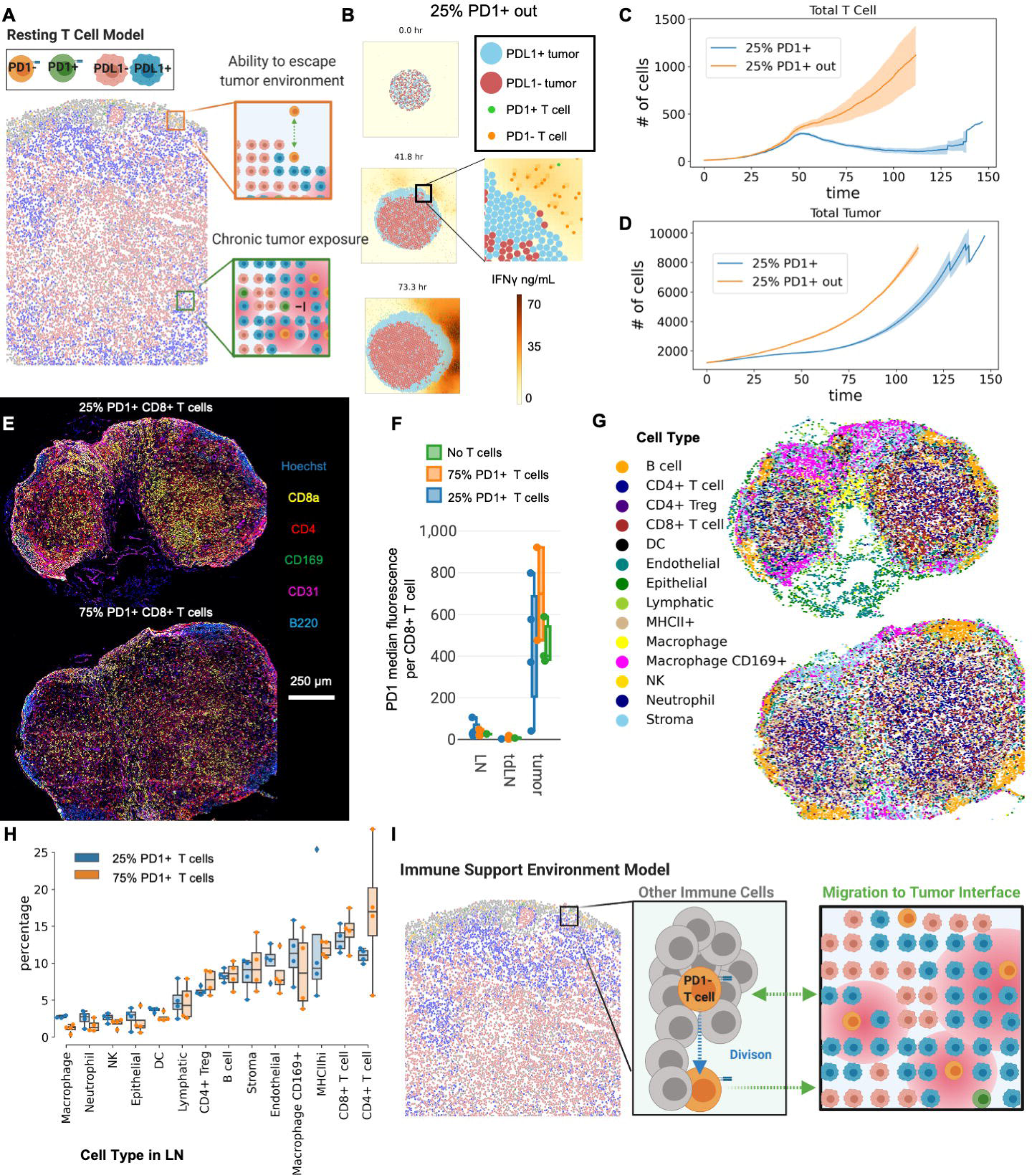
Conversion of tumor cell phenotype is more critical for tumor control than T cell phenotype preservation. **A)** Theoretical sketch of how T cells are able to escape the tumor microenvironment to promote long-term survival and ability to control the tumor. **B)** Snapshots of the tumor from agent-based modeling of a tumor treated with 25% PD1^+^ T cells that were initialized outside the tumor such that they can escape chronic exposure to tumor and limit exhaustion. **C)** Total number of T cells over time of simulation when T cells are initialized outside the tumor bed or inside the tumor (mean of n=4 replicates with shading showing SEM). **D)** Total number of tumor cells over time of simulation when T cells are initialized outside the tumor bed or inside the tumor (mean of n=4 replicates with shading showing SEM). **E)** Model of T cells supported by immune cells in a microenvironment within the tumor for preservation of phenotype, proliferation, killing, and tumor inhibition locally. **F)** Percentages of indicated adaptive immune cells in total cells in CODEX multiplexed images of tumors treated with 25% PD1^+^ T cells, 75% PD1^+^ T cells, or no T cells. **G)** Percentages of indicated innate immune cells in total cells from the CODEX multiplexed images from CODEX multiplexed images of tumors treated with 25% PD1^+^ T cells, 75% PD1^+^ T cells, or no T cells.

Interestingly, despite the much higher numbers of T cells located outside the tumor, in our simulations, the tumors with T cells initialized outside the tumor grew more quickly than tumors with T cells initialized inside (∼4500 cells vs. ∼2500 cells at 75 h) (Fig. 6D, Supplemental Fig. 7C-E). Consequently, delay of T cell exhaustion came at the cost of enhanced early tumor growth rates, which even larger numbers of T cells were not able to overcome in the long run. Since resting preserves phenotype, but does not enhance tumor outcome, a mechanism for enhanced tumor control in metabolically treated T cells is missing from our model.

We speculated therefore that T cells could come from outside the tumor enabling founder T cells to reside in supportive microenvironments like the lymph node, while daughter cells migrate into the tumor. To test this hypothesis, we harvested lymph nodes from mice with tumors treated with 75% PD1+ T cells, 25% PD1+ T cells, or No T cells treated 3 days before. We then created a tissue array of all these lymph nodes and imaged them simultaneously with CODEX multiplexed imaging (Fig. 6E). Comparing the median marker expression of PD1 for CD8+ T cells found in the lymph node to the CD8+ T cells in the tumor showed increased levels of PD1 in the tumor (Fig. 6F). This agrees with our hypothesis that founder T cells reside in protected lymph node environments sending daughter cells to tumors. However, we did not observe drastic differences of PD1 expression between treatment conditions and did not explain the discrepancy between treatments.

To see if the percentage of CD8+ T cells were different between the two conditions we segmented and clustered cell types in the CODEX lymph node datasets (Fig. 6G). This analysis showed that there were no differences in cell type percentages within the lymph nodes of both treatments (Fig. 6H). Since we did not observe differences between conditions within the lymph node, perhaps this concept of supportive microenvironments for T cells extends to the tumor (Fig. 6I). We propose that T cells actively build microenvironments within the tumor to support T cell phenotype and function. This led us to perform extensive studies evaluating the tumor microenvironment composition surrounding T cells and indicates immune cell supportive microenvironments provide critical support for productive T cell killing (*Hickey et. al., cosubmitted*).

## DISCUSSION

We developed a scalable agent-based model of T cell therapy of tumors to complement, leverage, and probe CODEX multiplexed imaging datasets of T cell-treated tumors. We specifically explored the importance of tumor phenotype on T cell therapy efficacy by incorporating molecular switches of cellular phenotype and function that led to tissue-level phenomena in our data-informed multiscale agent-based model. We then used findings from the simulations to guide design of an *in vivo* experiment and choice of antibodies for use in CODEX imaging to catalog cell types and transitions we saw within our models. This synergistic back-and-forth of model and data across biological scales revealed critical design components of effective T cell therapies (Fig. 7).

**Figure 7:**
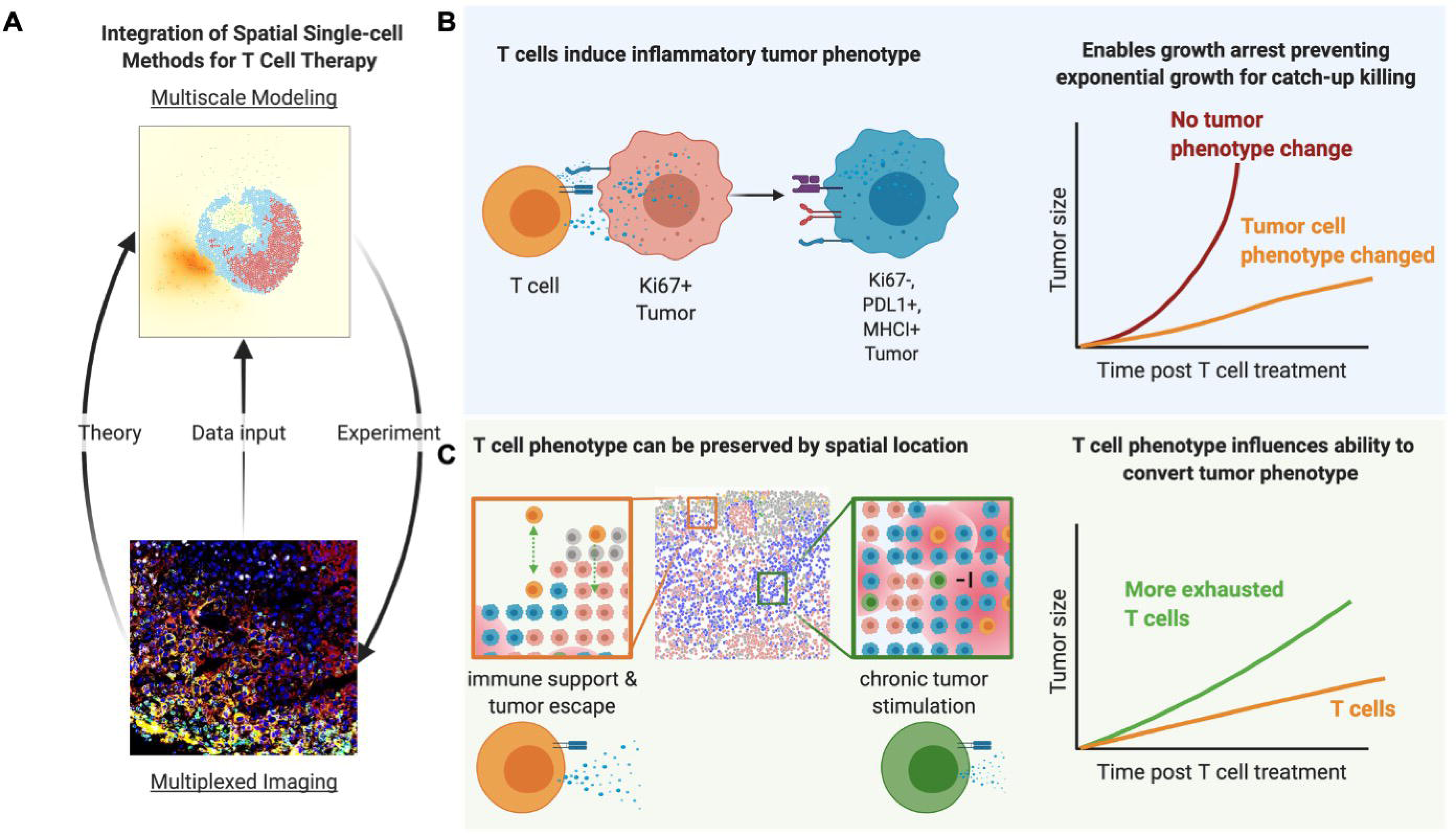
Integrating Multiplexed Imaging and Multiscale Modeling Identifies Tumor Phenotype Transformation as a Critical Component of Therapeutic T Cell Efficacy. **A)** Implementing both multiplexed imaging and multiscale models simultaneously allowed us to exploit a theory-experiment cycle that both enriched hypotheses for targeted experiments and improved model design. We reconstructed the complexity of the tumor microenvironment across multiple biological scales by creating data-informed multiscale agent-based models. We incorporated molecular switches of cellular phenotype and function that led to tissue-level phenomena. We also deconstructed the complexity of the tumor microenvironment across multiple biological scales by using CODEX to image 42 molecular markers quantified at the single-cell level. **B)** T cell phenotype also impacts its ability to change tumor phenotype, and a focus on converting tumor phenotype results in greater control than minimizing T cell exhaustion. **C)** Spatial analysis of CODEX and *in silico* experiments demonstrated that the spatial positioning of T cells influenced T cell phenotype.

The results indicated that tumor phenotype considerably influences the ability of T cells to control tumor cell growth through inhibiting proliferation and increasing killing. Most recent work has focused on controlling T cell phenotype for both the secretion of killing molecules and self-preservation (Blank et al., 2019; Gattinoni et al., 2017). We found that T cell phenotype also impacts its ability to change tumor phenotype, and a focus on converting tumor phenotype results in greater control than minimizing T cell exhaustion.

Comparing *in silico* results to CODEX multiplexed imaging reinforced the importance of T cell phenotype and influence on tumor phenotype; however, CODEX data indicated that T cells were able to change tumor phenotype and minimize T cell exhaustion. Most methods of analysis of T cell phenotype preservation are focused on molecular mechanisms of control since most assays require dissociation of tumors or *ex vivo* manipulation of T cells (Galletti et al., 2020; Krishna et al., 2020). In contrast, both multiscale modeling and multiplexed imaging preserve spatial features of the data, and our spatial analysis of CODEX and *in silico* experiments demonstrated that the spatial positioning of T cells influenced T cell phenotype. In the comparison to CODEX data, this suggested that our model was missing a mechanism for T cell phenotype preservation.

This incongruence motivated parallel research on T cell therapies, where we observed that therapeutic T cell phenotype changes the structure and cellular composition of the tumor microenvironment (*Hickey et. al., cosubmitted*). We found in this other work that T cells create distinct multicellular neighborhoods based on their phenotype and molecular expression profiles. For example, 2HC T cell treated tumors result in more productive T cell and tumor neighborhoods, whereas T cells not treated with 2HC secrete more anti-inflammatory cytokines and have T cell and tumor areas also enriched with regulatory neighborhoods. Thus, T cells should be engineered with the ability to transform tumor cell phenotype and to be agents of structural change of the tumor microenvironment.

In future work, driven by molecular data acquired through multiplexed tissue imaging, the model should be expanded to include other cell types and anti-inflammatory molecules. Since the *Vivarium* framework for multiscale modeling is modular and compositional, it will be straightforward to add additional cell types with different representations of internal mechanisms and environmental interactions. Not straightforward are the selection of molecular features and phenotypes that will increase the accuracy of the model under a broader range of conditions. Although we have some clues about missing components from our CODEX and RNA dataset (*Hickey et. al., cosubmitted*), additional data collection will be needed. For example, single-cell RNA sequencing of the cell types within the tumor will shed light on key molecular and cellular interactions. Building this complexity within *in silico* models will be necessary because as the number of intercellular connections are increased, it will become more difficult to recapitulate and deconvolute these networks within *in vitro* systems.

The modularity of the model will enable our group and others to build from this starting point to investigate the effects of various T cell-based therapies in solid tumors. Integration of models and measurements across biological scales with spatial features preserved will enable decoding of the rules that govern complex networks from biological and clinical samples. We and others can leverage this model as a template for integrating and using the growing number of spatial datasets, like CODEX imaging datasets, in other disease settings (Consortium, 2019; Hickey et al., 2021c; Rozenblatt-Rosen et al., 2020). Integration of multiscale modeling and imaging data will enable better interpretation through leave-one-out experiments and ensemble simulations while simultaneously increasing the complexity and accuracy of agent-based models. Finally, the ability to simultaneously evaluate interactions across scales will guide development of better therapies that interrupt problematic networks or create beneficial ones (Baertsch et al., 2021).

## ACKNOWLEDGEMENTS

This work was supported by the U.S. National Institutes of Health (2U19AI057229-16, 5P01HL10879707, 5R01GM10983604, 5R33CA18365403, 5U01AI101984-07, 5UH2AR06767604, 5R01CA19665703, 5U54CA20997103, 5F99CA212231-02, 1F32CA233203-01, 5U01AI140498-02, 1U54HG010426-01, 5U19AI100627-07, 1R01HL120724-01A1, R33CA183692, R01HL128173-04, 5P01AI131374-02, 5UG3DK114937-02, 1U19AI135976-01, IDIQ17X149, 1U2CCA233238-01, 1U2CCA233195-01), the U.S. Department of Defense (W81XWH-14-1-0180, W81XWH-12-1-0591), the U.S. Food and Drug Administration (HHSF223201610018C, DSTL/AGR/00980/01), Cancer Research UK (C27165/A29073), the Bill and Melinda Gates Foundation (OPP1113682), the Cancer Research Institute, Hope Realized Medical Foundation (209477), the Silicon Valley Community Foundation (2017-175329 and 2017-177799-5022), and the Rachford & Carlotta A. Harris Endowed Chair to GPN. JWH was supported by an NIH T32 Fellowship (T32CA196585) and an American Cancer Society - Roaring Fork Valley Postdoctoral Fellowship (PF-20-032-01-CSM). EA was supported by an NIH F32 Fellowship from NIGMS (F32GM137464) and by the Paul G. Allen Frontiers Group via an Allen Discovery Center at Stanford. The authors thank Ryan Spangler for helping set up the simulation development environment. Some figures were created with BioRender.com.

## AUTHOR CONTRIBUTIONS

JWH and EA conceived and developed the agent-based model, analyzed data, and wrote the manuscript. JWH and NH completed *in vitro* and *in vivo* based experiments with T cells. ML researched parameters for the model system. JS, MC, and GPN supervised the project, provided resources and feedback, and helped in writing the manuscript.

## DECLARATION OF INTERESTS

GPN has equity in and is a scientific advisory board member of Akoya Biosciences, Inc. The other authors declare no competing interests.

## Model Development

### The *Vivarium* framework

*Vivarium* is an open-source software tool for multi-scale modeling. The aim was to make it easier for scientists to define any imaginable mechanistic model, combine it with existing models, and execute them together as an integrated simulation. It provides an interface that makes individual simulation tools into modules that can be wired together, parallelized across multiple CPUs, and simulated across many spatial and temporal scales (Agmon et al., 2021).

*Vivarium’s* basic elements are processes and stores (Fig. 2C). A *Vivarium process* is an object that contains parameters and the update function, which describes the inter-dependencies between the variables and how they map from one time to the next. A *store* is an object that holds the system’s state variables and applies the processes’ updates. Processes include *ports*, which allow users to wire processes together through variables in shared stores with connections called a *topology*. Multiple processes can be wired together as integrated models called *composites*. These models are implemented in a nested hierarchy, which has stores within stores to allow an environmental model to run at the top of the hierarchy, with individual agents running in parallel within the model.

### Cell Processes

The model is composed of two major cell types (T cells and tumor cells), each with two separate phenotypes. Each cell type has an associated *Vivarium* process that represents the mechanisms that make a cell switch between phenotypes. These processes define fundamental rules that govern cellular interactions with the other cell types and with the inputs it receives from the environment. The tumor process is focused on two phenotypic states: proliferative with low levels of immune molecules (MHCI and PDL1) and quiescent with high levels of immune molecules (MHCI and PDL1). Its transition from the proliferative state is dependent on the level of IFNγ secreted by T cells. Both tumor types can be killed by receiving cytotoxic packets from the T cells. The T cell process is focused on two phenotypic states. The PD1^-^ T cells secrete larger amounts of immune molecules (IFNγ and cytotoxic packets) than PD1^+^ T cells. These immune molecules impact the state and death of tumor cells. The transition from the PD1^-^ state to the PD1^+^ state is dependent on the length of time the T cell is engaged with tumor cells. Each process was tested individually to meet expected outcomes based on literature or lab data. Testing the processes individually reveals whether underlying parameters derived from literature values or primary data accurately represent behavior expected based on such research.

### Cell Composites

The T cell and tumor processes are combined with additional processes to create T cell and tumor composite agents. These include a division process, which waits for division to be triggered and then carries out division; a death process, which waits for death to be triggered and then removes the agent; and a local field process, which interfaces the external environment to support uptake and secretion for each agent. Testing individual composite cells adds additional complexity and is another accuracy check of the model.

### Tumor Microenvironment

The Tumor Microenvironment is a composite model that simulates a 2D environment with agents that can move around in space, exchange molecules with their neighbors, and exchange molecules with a molecular field. A *neighbors* process models individual agents as circular rigid bodies that can move, grow, and collide. This process tracks the locations of individual agents and handles the exchanges between neighboring cells. A *diffusion* process operates on the molecular fields of IFNγ, and handles the cells uptake and secretion from the environment.

### Connecting Cell Composites in the Tumor Microenvironment

After validating all individual processes, we connected processes and composites and endowed individual elements with additional behaviors like migration. In our model, the T cells can interact with tumor cells through 1) the TCR on T cells and MHCI molecules on tumor cells to activate T cells, induce IFNγ and cytotoxic packet secretion, and inhibit T cell migration, 2) PD1 receptor on T cells and PDL1 receptor on tumor cells that can inhibit T cell activation and induce apoptosis, and 3) indirectly through secretion of IFNγ by T cells, which is taken up by tumor cells to cause a state switch to upregulate MHCI and PDL1 and decrease proliferation.

For additional information on model development see our documented code base and README at https://github.com/vivarium-collective/tumor-tcell.

## T Cell Culture and Stimulation

### Mice

B6 and PMEL transgenic mice were maintained per guidelines approved by Stanford University’s Institutional Review Board. C57BL/6J and PMEL mice were purchased from Jackson Laboratories.

### Immune cell isolation

Murine cells were obtained from adult mouse lymph nodes and spleens. Obtained cells were treated with ACK lysis buffer to lyse red blood cells, and lysates were filtered through cell strainers to isolate splenocytes.

### T cell media

Supplemented media was made with RPMI 1640 media with glutamine, 1x non-essential amino acids, 1 mM sodium pyruvate, 0.4x vitamin solution, 92 µM 2-mercaptoethanol, 10 µM ciprofloxacin, and 10% fetal bovine serum.

### CD3-coated plate preparation

To each well of a 96-well, U-bottomed plate was added 50 µL of a solution of 5 µg/mL anti-CD3 (Bioxcell, clone 145-2C11) in PBS. After incubation at 4 °C overnight, liquid was decanted.

### T cell stimulation

Isolated murine immune cells were stimulated by incubation with 1 μM cognate peptide GP100 (KVPRNQDWL) and 50 IU/mL IL-2. For 2-hyroxycitrate (2HC) conditions, 2HC was added to culture media at a concentration of 5 mM. Cells were seeded at a density of 2-5×10^6^ cells/mL. Cells were fed with additional IL-2 in T cell media every other day. On day 5, cells were added to CD3-coated plates in culture media containing 2 µg/mL anti-CD28 (Bioxcell, clone 37.51). On day 8 cells were removed from plates and plated on uncoated plates and fed with IL-2-containing media until day 10.

## *In Vitro* T Cell Killing Assay

T cell killing and tumor cell growth rates were determined using the xCELLigence Real-Time Cell Analysis platform. Wells of xCELLigence E-plates were coated with gold nanoparticles and electrical potential was passed across the plate every 15 min. Monitoring of changes in the electrical impedance enabled quantification of adherent cells over time. For these assays, T cells were expanded. B16-F10 melanoma cells were split and then left as control cells or treated with 10 ng/mL IFNγ for 24 hours prior to plating. After the pretreatment, 10,000 B16-F10 cells were plated in each well of the xCELLigence E-plate and allowed to adhere for 12 h. Next, T cells were added at 1:1 effector to target ratio. The growth of tumor cells and killing by T cells was monitored for up to 24 h. Killing was calculated by normalizing the cell index of each well to the time point just before addition of the T cells and then quantifying the differences between T cell and control wells without T cells over time.

## CyTOF Phenotyping

### Antibodies

Primary antibody transition metal-conjugates were prepared in-house using 100-μg antibody lots and the MaxPAR antibody conjugation kit (DVS Sciences) according to the manufacturer’s recommended protocol. Following conjugation, antibodies were diluted in Candor PBS Antibody Stabilization solution and stored at 4 °C. *For additional details of the antibody panel see (*Hickey et. al., cosubmitted*).

### Staining

Following T cell stimulation on day 10, 2×10^6^ cells were stained with cisplatin at 25 μM in PBS in 1 mL for 1 min at 4° C, quenched with 1 mL of fetal bovine serum, and washed with cell staining medium (CSM; PBS with 0.5% bovine serum albumin and 0.02% sodium azide). Cells were blocked with FcBlock (0.25 μg/1×10^6^ cells) for 15 min at room temperature, then the surface antibody cocktail was added and incubated 1 h at room temperature on a shaker at 100 rpm. Cells were washed with CSM and then with PBS. Cells were fixed and stained with intercalators overnight at 4° C in a solution of 1.6% PFA in PBS. The next day, the cells were washed once with CSM and twice with doubly distilled water, resuspended in doubly distilled water, and analyzed using CyTOF.

## Flow Cytometry for Intratumoral T Cell Measurement

### In vivo tumor model

On day 0, B6 mice were injected with 2×10^5^ B16-F10 melanoma tumor cells. On day 0, immune cells were isolated from a PMEL mouse and stimulated as described above for 10 days to produce stimulated T cells for adoptive transfer. On day 9, mice were given a central dose of 500 cGy, which induces transient lymphopenia (Wrzesinski et al., 2010). On day 10, T cells cultured *ex vivo* were harvested and adoptively transferred intravenously in volumes of 100 µL with 1×10^6^ stimulated T cells per mouse. Tumors were harvested 3 days after treatment with stimulated T cells.

### Flow cytometry staining

Harvested tumors were dissociated by maceration over a sterile 70-µm cell strainer with frequent washes of PBS. Cells were then stained with a mixture of a 1:100 PBS solution of APC-conjugated rat anti-mouse CD8a, clone 53–6.7 (BD Pharmingen), PerCP-conjugated rat anti-mouse CD45, clone 30-F11 (Biolegend), and 1:1000 of LIVE/DEAD Fixable Violet Dead Cell Stain (ThermoFisher) for 15 min at 4 °C. Cells were then washed with FACS wash buffer and analyzed using a BD FACSCalibur flow cytometer with CellEngine gating for live cells.

## Neighborhood Analysis of Simulation Output

Neighborhood analysis was performed as described previously (Schürch et al., 2020) on simulation output for day 3 after treatment of tumors with T cells of different phenotype compositions. Briefly a window size of 10 nearest neighbors was taken across the tissue cell type maps clustered into 5 neighborhoods. These clusters were mapped back to the tissue and evaluated for cell type enrichments to determine overall structure.

## DATA AND CODE AVAILABILITY

All of the code can be found on the repository site https://github.com/vivarium-collective/tumor-tcell. The README file documents how this can be used as a Python library or cloning the repository locally. Data from the CODEX experiments and initializations can also be found within the repository in *data*.

## SUPPLEMENTAL MATERIAL

Supplemental figures, tables, and legends are included in the attached pdf. Supplemental videos 1-3 are included separately as mp4 files with following descriptions below.

**Supplemental Video 1:** Time lapse of simulation of tumor microenvironment treated with 25% PD1^+^ T cells.

**Supplemental Video 2:** Time lapse of simulation of tumor microenvironment treated with 75% PD1^+^ T cells.

**Supplemental Video 3:** Time lapse of simulation of tumor microenvironment treated with no T cells.

