## Supplementary Material for "Integrating Multiplexed Imaging and Multiscale Modeling Identifies Tumor Phenotype Transformation as a Critical Component of Therapeutic T Cell Efficacy"

SUPPLEMENTAL FIGURES 1-7

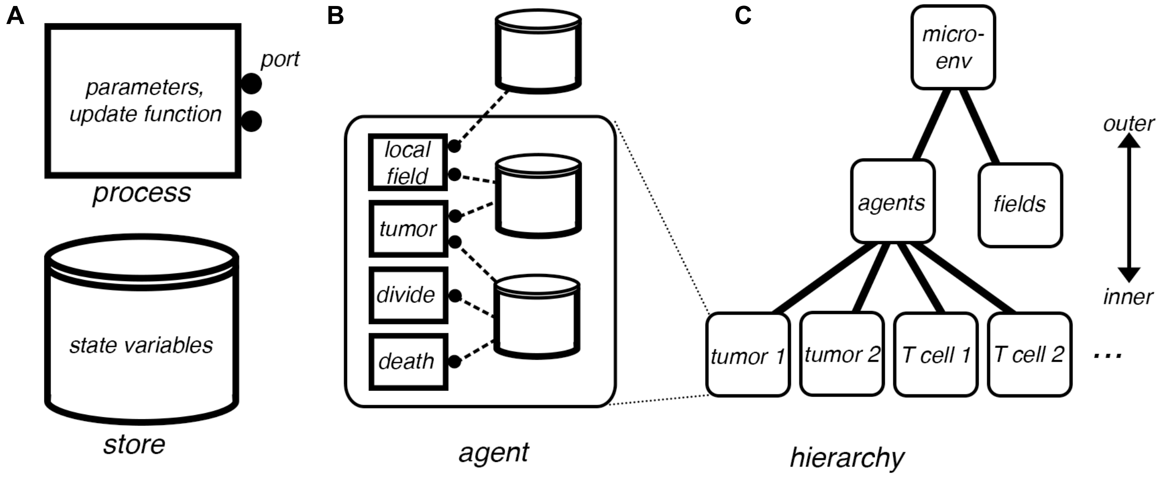


**Supplemental Figure 1:** Schematic of the tumor/T cell microenvironment model's Vivarium interface, illustrating the simulation's formal structure. **A)** *Vivarium's* basic elements. The *process*, shown as a rectangular flowchart symbol, is a modular model that contains the parameters, an update function, and ports. The *store*, shown as the flowchart symbol for a database, holds the state variables and schemas that determine how updates are handled. **B)** *Agents* are bundles of *processes* and *stores* wired together (called *composites* in *Vivarium* terminology) across a single level. The figure shows the main tumor process wired with processes called local field (for reading the external concentrations), divide (for dividing the cell), and death (for removing the cell from the simulation). **C)** Cell agents are *compartments* embedded in a hierarchy depicted here as a hierarchical network with discrete layers. Outer *compartments* are shown above and inner compartments below. The cells all exist under "agents" and the external fields are held in an adjacent *store* called "fields".

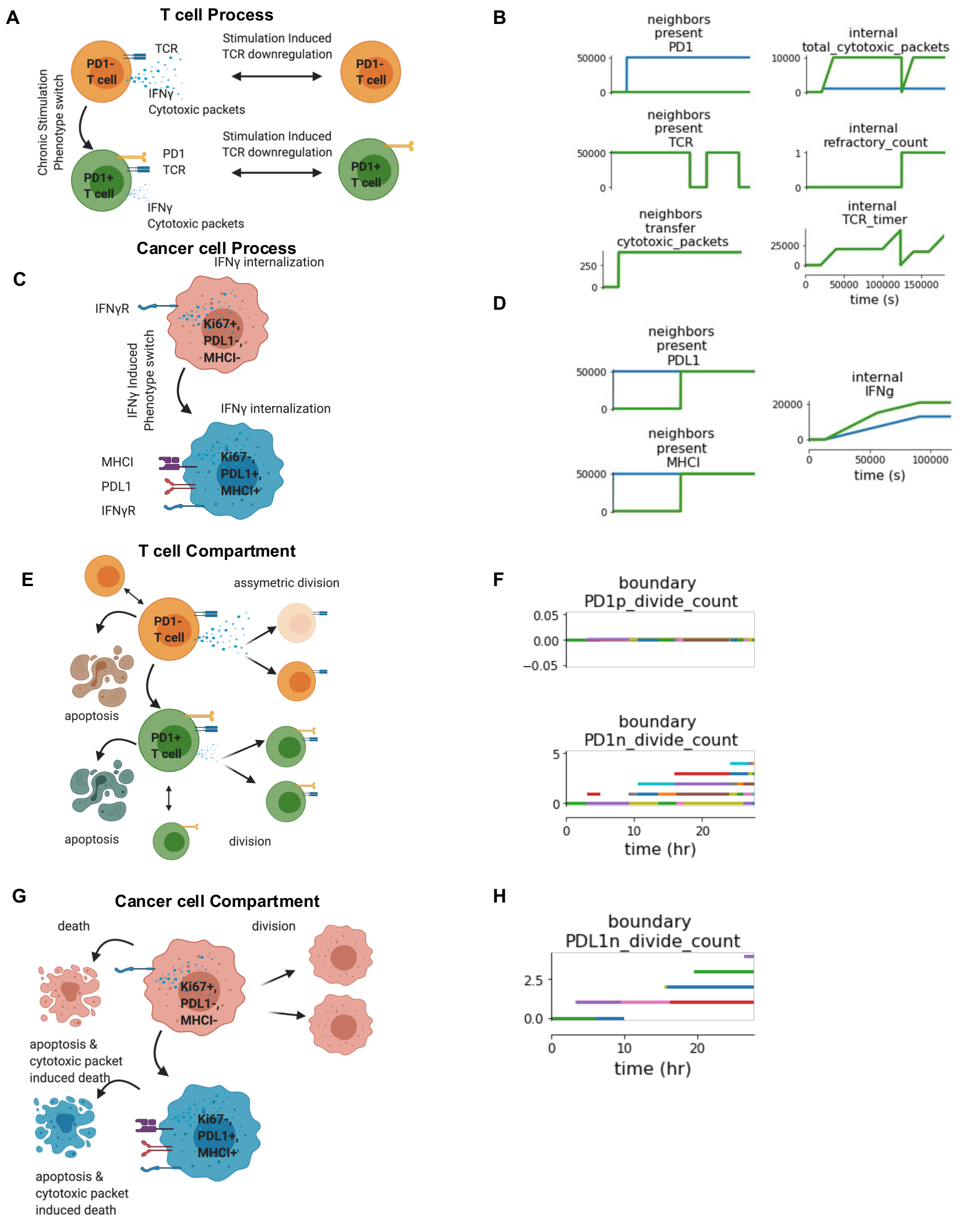


**Supplemental Figure 2:** Representation and development of individual components of the multiscale agent-based model. **A)** T cell process is composed of two T cell states. PD1⁻ CD8⁺ T cells become PD1⁺ T cells upon chronic stimulation and both PD1⁺ T cells and PD1⁻ T cells can downregulate TCR. Both T cell types express TCR, IFNγ, and produce cytotoxic packets. Molecular regulation is governed by activation and stimulation of tumor cells. **B)** Representative output of simulating only the T cell process. **C)** Tumor cell process is composed of two tumor cell states. $Ki67^+$ $PDL1^-$ $MHCI^-$ tumor cells can become $Ki67^-$ PDL1, $MHCI^+$ tumor cells upon exposure to IFNγ,. Both tumor cell types express IFNγR. Molecular regulation is governed by interaction with T cells. **D)** Representative output of simulating only the tumor cell process. **E)** The T cell compartment extends the T cell process by adding division and death processes that can be asymmetric. **F)** Representative output of simulating the T cell compartment. **G)** The tumor cell compartment adds proliferation and death processes. **H)** Representative output of simulating the tumor cell compartment.

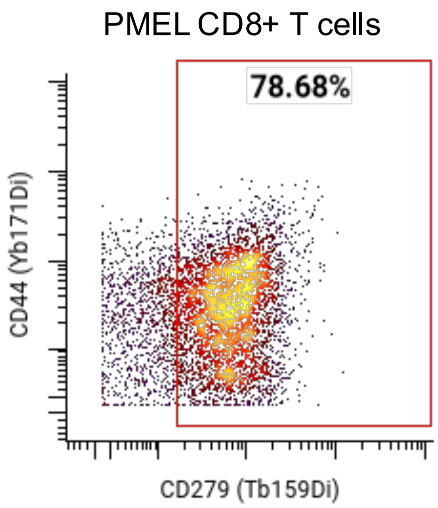


**Supplemental Figure 3:** Conditions used to initialize the T cell phenotype in the multiscale model. Nearly 80% of restimulated T cells used for the *in vitro* killing assay express PD1 (CD279) as plotted versus CD44 as measured by CyTOF. This condition was used to initialize the T cell phenotype for the agent-based model.

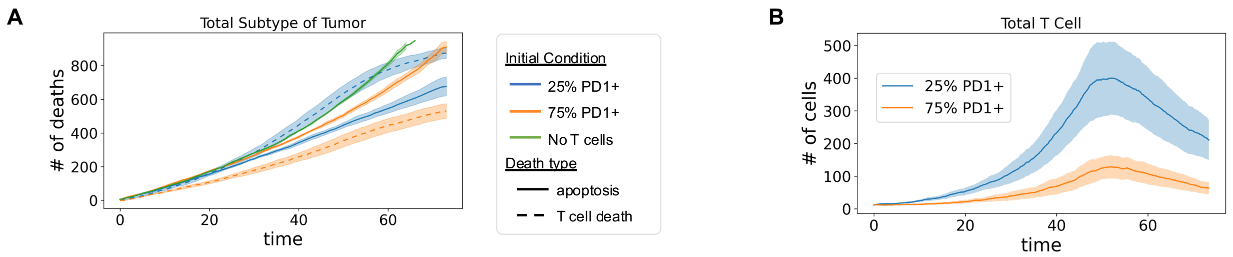


**Supplemental Figure 4:** Agent-based modeling results of treatment with 25% PD1$^+$ T cells, 75% PD1$^+$ T cells, or no T cells. **A)** Total number of tumor cell deaths quantified for T cell-induced killing and apoptotic events over 60-h simulation of treatment with 25% PD1$^+$ T cells, 75% PD1$^+$ T cells, or no T cells (mean of n=4 replicates with shading showing SEM). **B)** Number of T cells during 60-h simulation of treatment with 25% PD1$^+$ and 75% PD1$^+$ T cells (mean of n=4 replicates with shading showing SEM).

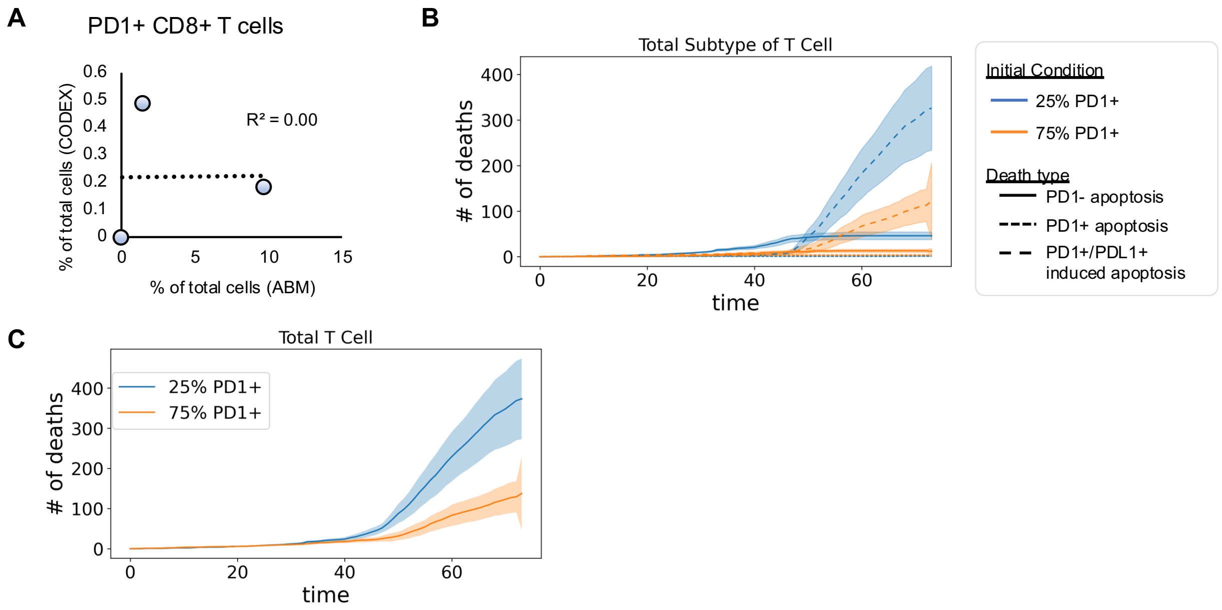


**Supplemental Figure 5:** Differences in preservation of T cell phenotype between CODEX multiplexed imaging and multiscale agent-based modeling highlights potential mechanism for phenotype preservation. **A)** Correlation plots of percent of $PD1^+$ $CD8^+$ T cells remaining after 3 days of T cell therapy for CODEX multiplexed imaging of *in vivo* experiments versus *in silico* simulations. **B)** Total number of T cell deaths grouped by type in tumors treated with 25% $PD1^+$ T cells or 75% $PD1^+$ T cells (mean of n=4 replicates with shading showing SEM). **C)** Number of total T cell deaths in tumors treated with 25% $PD1^+$ T cells or 75% $PD1^+$ T cells (mean of n=4 replicates with shading showing SEM).

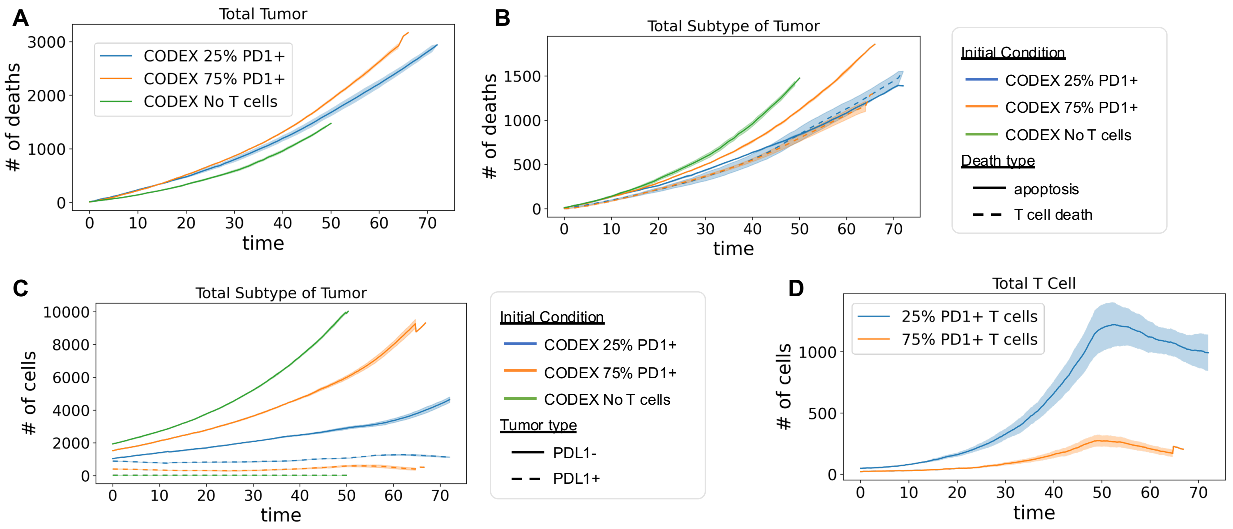


**Supplemental Figure 6:** Results from agent-based modeling simulations initialized with CODEX multiplexed imaging data. **A)** Total number of tumor cell deaths in simulations of treatment with 25% PD1$^+$ T cells, 75% PD1$^+$ T cells, or no T cells. B) Tumor cell deaths due to T cell-induced killing and apoptotic events in simulations of indicated treatments. **C)** Numbers of PDL1$^+$ and PDL1$^-$ tumor cells for each of the three treatment groups across the simulation. **D)** Number of T cells across the duration of the agent-based model for 25% PD1$^+$ and 75% PD1$^+$ T cell treatment conditions. For all panels: mean of n=4 replicates with shading showing SEM.

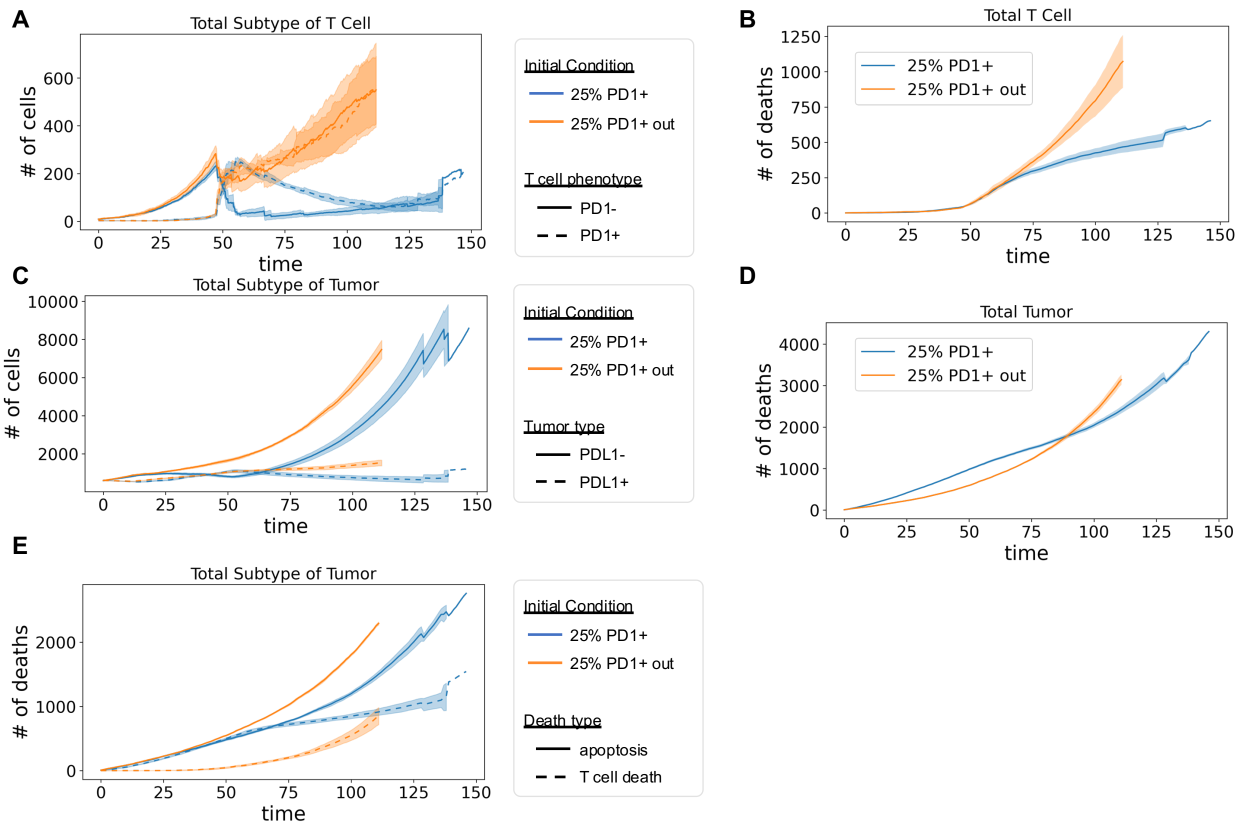


**Supplemental Figure 7:** Initialization of therapeutic T cells outside the tumor bed allows T cell phenotype preservation and greater numbers of T cells but faster tumor growth. **A)** Number of T cells by phenotype (PD1$^+$ and PD1$^-$) quantified across the duration of the agent-based model for 25% PD1$^+$ initialized outside and 25% PD1$^+$ T cell initialized inside conditions. **B)** Total number of T cell deaths quantified for 25% PD1$^+$ initialized outside and 25% PD1$^+$ T cell initialized inside conditions. **C)** Total number of PDL1$^+$ and PDL1$^-$ tumor cells for 25% PD1$^+$ initialized outside and 25% PD1$^+$ T cell initialized inside conditions. **D)** Total number of tumor cell deaths for 25% PD1$^+$ initialized outside and 25% PD1$^+$ T cell initialized inside conditions. **E)** Total number of tumor cells deaths by T cell-induced killing and apoptotic events by tumor cell phenotypes in simulations with 25% PD1$^+$ initialized outside and 25% PD1$^+$ T cell initialized inside conditions. In all panels: mean of n=4 replicates with shading showing SEM.

SUPPLEMENTAL TABLES

Supplemental Table 1: Key model parameters and the source from the literature.

| Model name | Value | Source | Location |
|---|---|---|---|
| refractory_count_threshold | 3 | (Zhao et al., 2020) | t_cell.py |
| PD1n_divide_threshold | 5 | | |
| activation_time | 21600 s | (Gallegos et al., 2016; Salerno et al., 2017) | |
| activation_refractory_time | 43200 s | | |
| death_PD1p_14hr | 0.35 | (Petrovas et al., 2007) | |
| death_PD1n_14hr | 0.1 | | |
| death_PD1p_next_to_PDL1p_14hr | 0.475 | (Dong et al., 2002; Tang et al., 2015) | |
| PD1n_IFNg_production | 1.62e4 molecules/cell/s | (Bouchnita et al., 2017; Zelinskyy et al., 2005) | |
| PD1p_IFNg_production | 1.62e3 molecules/cell/s | | |
| PD1n_growth_28hr | 0.9 | (Petrovas et al., 2007; Vodnala et al., 2019) | |
| PD1p_growth_28hr | 0.2 | | |
| PD1n_migration | 10 μm/min | (Boissonnas et al., 2007) | |
| PD1p_migration | 5 μm/min | | |
| migration_MHCIp_tumor_dwell_velocity | 0 μm/min | (Thibaut et al., 2020) | |
| PD1n_migration_MHCIp_tumor_dwell_time | 25 min | | |
| PD1p_migration_MHCIp_tumor_dwell_time | 10 min | | |
| PD1n_migration_refractory_time | 35 min | | |
| PD1p_migration_refractory_time | 20 min | | |
| cytotoxic_packet_production | 40 /min | (Betts and Koup, 2004; Zhang et al., 2006) | |
| PD1n_cytotoxic_packets_max | 10000 | | |
| cytotoxic_transfer_rate | 400 /min | | |
| PD1p_cytotoxic_packets_max | 1000 | (Zelinskyy et al., 2005) | |
| MHCIn_reduction_production | 400 | (Böhm et al., 1998; Merritt et al., 2004) | |
| death_apoptosis | 0.5 | (Gong et al., 2017) | tumor.py |
| cytotoxic_packet_threshold | 128 | (Betts and Koup, 2004; Zhang et al., 2006) | |
| PDL1n_growth | 0.6 | (Eden et al., 2011) | |
| Max_IFNg_internalization | 21 molecules/min | (Celada and Schreiber, 1987) | |

| | | | |
|---|---|---|---|
| IFNg_threshold | 15000 molecules | (Hoekstra et al., 2020; Thibaut et al., 2020) | |
| reduction_IFNg_internalization | 2 | (El Darzi et al., 2017; Ersvaer et al., 2007) | |
| DIFFUSION_RATES: IFNg | 1.25e-3 $cm^2$/day | (Liao et al., 2014) | fields.py |
| decay: IFNg | 4.5 hr | (Kurzrock et al., 1985) | |
